# Inhibiting succinate release worsens cardiac reperfusion injury by enhancing mitochondrial reactive oxygen species generation

**DOI:** 10.1101/2022.04.27.489760

**Authors:** Alexander S. Milliken, Sergiy M. Nadtochiy, Paul S. Brookes

**Affiliations:** Department of Pharmacology and Physiology; Department of Anesthesiology and Perioperative Medicine, University of Rochester Medical Center

## Abstract

The metabolite succinate accumulates during cardiac ischemia. Within 5 min. of reperfusion, succinate returns to baseline levels via both its release from cells and oxidation by mitochondrial complex II (Cx-II). The latter drives reactive oxygen species (ROS) generation and subsequent opening of the mitochondrial permeability transition (PT) pore, leading to cell death. Targeting succinate dynamics (accumulation/oxidation/release) may be therapeutically beneficial in cardiac ischemia-reperfusion (IR) injury. It has been proposed that blocking monocarboxylate transporter 1 (MCT-1) may be beneficial in IR, by preventing succinate release and subsequent engagement of downstream inflammatory signaling pathways. In contrast, herein we hypothesized that blocking MCT-1 would retain succinate in cells, exacerbating ROS generation and IR injury. Using the mitochondrial ROS probe mitoSOX, and a custom-built murine heart perfusion rig built into a spectrofluorometer, we measured ROS generation *in-situ* during the first moments of reperfusion, and found that acute MCT-1 inhibition enhanced mitochondrial ROS generation at reperfusion, and worsened IR injury (recovery of function and infarct size). Both these effects were abrogated by tandem inhibition of Cx-II, suggesting that succinate retention worsens IR due to driving more mitochondrial ROS generation. Furthermore, using the PT pore inhibitor cyclosporin A, along with monitoring of PT pore opening via the mitochondrial membrane potential indicator TMRE, we herein provide evidence that ROS generation during early reperfusion is upstream of the PT pore, not downstream as proposed by others. In addition, pore opening was exacerbated by MCT-1 inhibition. Together, these findings highlight the importance of succinate dynamics and mitochondrial ROS generation, as key determinants of PT pore opening and IR injury outcomes.

## 1. INTRODUCTION

The metabolite succinate has a central role in tissue ischemia. Several mechanisms exist for ischemic succinate accumulation ^1, 2^, and this process is conserved across diverse tissues, species, and physiologic contexts ^3, 4^. However, rapid oxidation of accumulated succinate at the onset of tissue reperfusion is a key event in ischemia-reperfusion (IR) injury, the underlying pathology of myocardial infarction (heart attack) ^1, 2^. This has led to intense interest in succinate as a potential therapeutic target ^5-9^.

Approximately ⅓ of succinate accumulated during ischemia is rapidly oxidized by complex II (Cx-II) of the mitochondrial respiratory chain ^2, 10^. This leads to the generation of reactive oxygen species (ROS) via mechanisms that are thought to involve either reverse electron transport (RET) at complex-I (Cx-I) ^11-13^ or forward electron transport at complex-III (Cx-III) ^14-18^. Mitochondrial ROS generation potentiates opening of the mitochondrial permeability transition (PT) pore ^19-22^, which triggers necrotic cell death.

The remaining ⅔ of succinate accumulated during ischemia is released upon reperfusion, in a pH-dependent manner via monocarboxylate transporter 1 (MCT-1) ^10, 23^. The physiologic function of this released succinate is unclear, with suggestions that it may serve as a metabolic signal of hypoxia ^24^. Succinate is a ligand for the widely expressed succinate receptor (SUCNR1, formerly GPR91), which elicits a range of physiologic responses ^25^. Most relevant to IR injury, SUCNR1 signaling promotes inflammation via macrophage activation, which may contribute to the pathology of IR injury ^10, 26-30^. As such, it has been postulated that inhibiting MCT-1 would be beneficial in IR injury, via blunting of extracellular succinate → SUCNR1 signaling ^10, 31^.

In contrast, given the key role of intracellular succinate for ROS generation during reperfusion, we hypothesized herein that acute blockade of succinate release via MCT-1 would worsen IR injury, and that simultaneous blockade of Cx-II would abrogate this effect. This hypothesis was tested by measuring mitochondrial ROS generation *in-situ* in perfused mouse hearts using a custom-built perfusion rig within the chamber of a benchtop spectrofluorometer. In addition, since it has been proposed that PT pore opening itself may lie upstream of the burst of ROS seen upon reperfusion ^23, 32^, we used this apparatus to interrogate the temporal relationship between ROS generation and PT pore opening during early reperfusion.

## 2. MATERIALS AND METHODS

### 2.1 Animals and Reagents

Animal and experimental procedures complied with the National Institutes of Health *Guide for Care and Use of Laboratory Animals* (8^th^ edition, 2011) and were approved by the University of Rochester Committee on Animal Resources (protocol #2007-087). Male and female C57BL/6J adult mice (8-20 weeks old) were housed in a pathogen-free vivarium with 12 hr. light-dark cycles and food and water *ad libitum*. Mice were administered terminal anesthesia via intra-peritoneal 2,2,2-tribromoethanol (Avertin) ∼250 mg/kg. Avertin was prepared in amber glass vials and stored at 4°C for no more than 1 month. This agent was chosen for anesthesia since it does not impact cardioprotection as reported for volatile anesthetics or opioids ^33-35^, and does not have mitochondrial depressant effects as reported for barbiturates ^36^. Euthanasia occurred via cardiac extirpation (see below).

MitoSOX red was from Thermo (NJ, USA) and was stored aliquoted under argon prior to use. Fully oxidized mitoSOX was prepared by reaction with Fremy’s salt, as described elsewhere ^37^. AR-C155858 was from MedChemExpress (Monmouth Junction, NJ, USA). Unless otherwise stated, all other reagents were from Sigma (St. Louis MO, USA).

### 2.2 Perfused Mouse Hearts

Following establishment of anesthetic plane (toe-pinch response), beating mouse hearts were rapidly cannulated, excised and retrograde perfused at a constant flow (4 ml/min.) with Krebs-Henseleit buffer (KHB) consisting of (in mM): NaCl (118), KCl (4.7), MgSO_4_ (1.2), NaHCO_3_ (25), KH_2_PO_4_ (1.2), CaCl_2_ (2.5), glucose (5), pyruvate (0.2), lactate (1.2), and palmitate (0.1, conjugated 6:1 to bovine serum albumin). KHB was gassed with 95% O_2_ and 5% CO_2_ at 37°C. A water-filled balloon connected to a pressure transducer was inserted into the left ventricle and expanded to provide a diastolic pressure of 6-8 mmHg. Cardiac function was recorded digitally at 1 kHz (Dataq, Akron OH) for the duration of the protocol. Following equilibration (10-20 min.) ischemia-reperfusion (IR) injury comprised 25 min. global no-flow ischemia plus 60 min. reperfusion. Hearts were then sliced and stained with triphenyltetrazolium chloride (TTC) for infarct quantitation by planimetry (red = live tissue, white = infarct). Infarction analysis was blinded to experimentalists.

The following conditions (Figure 1F) were examined: **(i) Control:** DMSO vehicle infusion 5 min. prior to ischemia and 5 min. into reperfusion. **(ii) S1QEL:** 1.6 µM S1QEL1.1, delivered 5 min. prior to ischemia and 5 min. into reperfusion. **(iii) DMM5:** 5 mM dimethyl malonate at the onset of reperfusion and 5 min. into reperfusion. **(iv) AR:** 10 µM AR-C155858 infusion, 5 min. prior to ischemia and 5 min. into reperfusion. **(v) AR + DMM5:** 10 µM AR-C155858 infusion, 5 min. prior to ischemia and 5 min. into reperfusion plus 5 mM dimethyl malonate at the onset of reperfusion and 5 min. into reperfusion. **(vi) AR + DMM10:** 10 µM AR-C155858 infusion, 5 min. prior to ischemia and 5 min. into reperfusion plus 10 mM dimethyl malonate at the onset of reperfusion and 5 min. into reperfusion. **(vii) AR + AA5:** 10 µM AR-C155858 infusion, 5 min. prior to ischemia and 5 min. into reperfusion plus 100 nM atpenin A5 at the onset of reperfusion and 5 min. into reperfusion. **(viii) CsA:** 0.8 µM cyclosporin A infusion ^38^, 5 min. prior to ischemia and 5 min. into reperfusion. Simultaneously with these conditions, hearts were delivered either 1.5 µM mitoSOX for 5 min. or 500 nM tetramethylrhodamine ethyl ester (TMRE) for 20 min., prior to ischemia.

**Figure 1.**
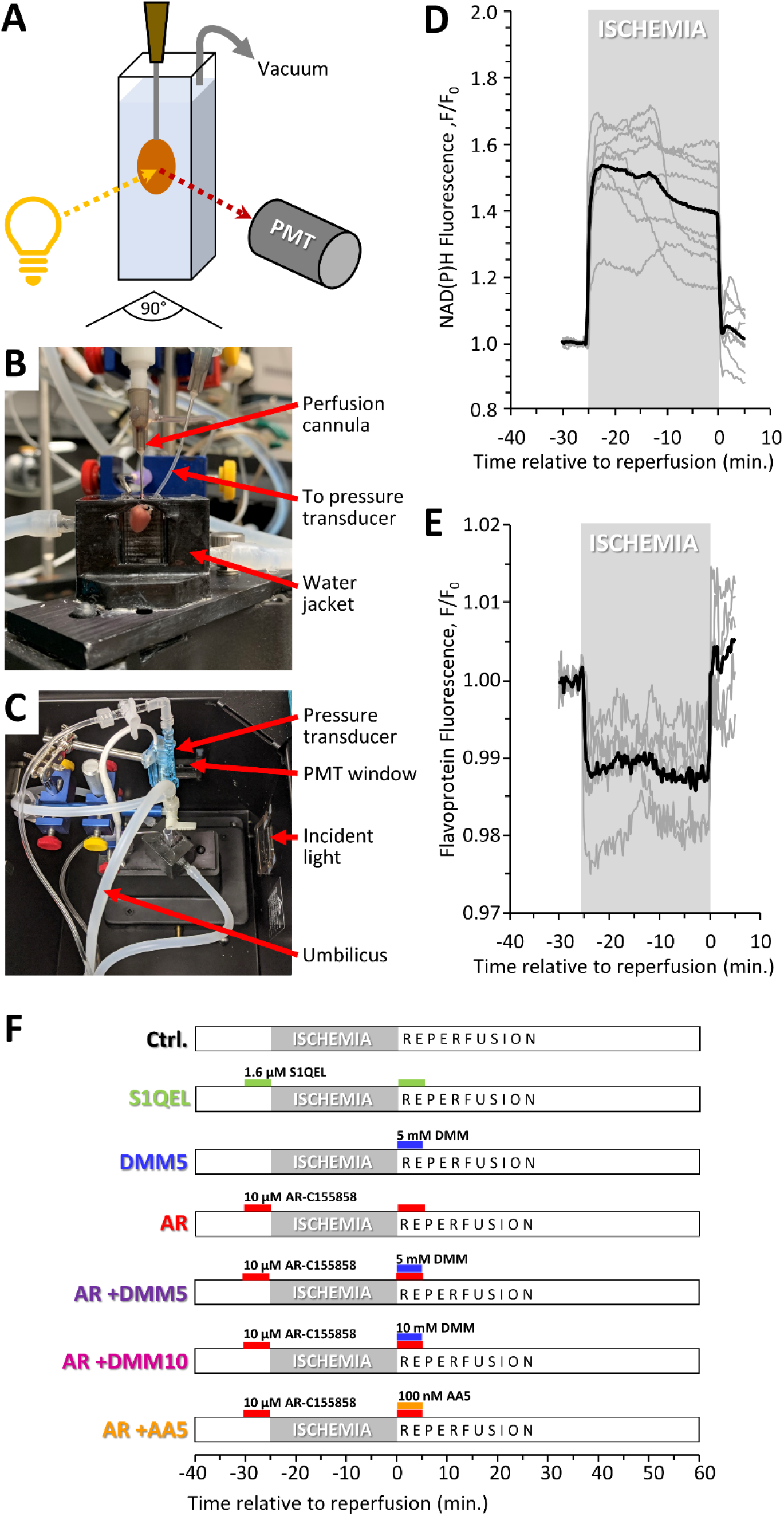
Fluorescence Cardiac Perfusion System. **(A):** Schematic showing arrangement of the perfused mouse heart relative to light source and photomultiplier tube (PMT). **(B):** Photograph of heart in-situ prior to mounting inside spectrofluorometer. **(C):** Photograph of cardiac perfusion apparatus from above mounted inside spectrofluorometer. Components of the system are labeled, including a custom 3D-printed water-jacketed cuvet holder, heated umbilical lines to deliver KHB at 37°C to the heart, and vacuum line to remove perfusate/effluent. **(D):** NAD(P)H autofluorescence (340/460 nm) during ischemia and reperfusion. Traces in gray are individual hearts, with the average (N=7) shown in black. **(E):** Flavoprotein autofluorescence (462/520 nm) during ischemia and reperfusion. Traces in gray are individual hearts, with the average (N=4) shown in black. Fluorescence data are normalized to pre-ischemic levels. **(F):** Schematic showing design of the 7 experimental perfusion conditions examined herein. Where indicated, S1QEL1.1 (Cx-I ROS inhibitor), dimethylmalonate (Cx-II inhibitor), AR-C155858 (MCT-1 inhibitor) or atpenin A5 (Cx-II inhibitor) were administered at the listed concentrations. Colors and terms used to denote each condition are used throughout the remainder of the Figures.

### 2.3 Perfusion Apparatus Within Spectrofluorometer

A cardiac perfusion apparatus was custom-built with an umbilicum, to position the heart within the light-proof enclosure of a Varian/Cary Eclipse benchtop spectrofluorometer (Agilent, Santa Clara CA, USA), as shown in Figure 1A-1C. Hearts were placed against the wall of a water-jacketed cuvette maintained at 37°C in which the excitation light source strikes the left ventricle of the heart at a 45° angle relative to the photomultiplier tube (PMT) window. The cuvet holder was 3D printed (.stl file in supplementary materials). Data was collected using Cary WinUV Kinetics software, which permitted real-time fluorescence monitoring (1 s. reads / 15 s. cycle, PMT voltage 600, Ex/Em slit width = 5 nm) for the duration of the perfusion protocol. At the heart rates observed (438 ± 54 bpm, mean ± SD, N=54) a 1 s. fluorescent read averages across ∼7 heart beats, so gating for motion artifacts was unnecessary. Initial validation was performed by monitoring endogenous NAD(P)H fluorescence (λ_EX_ 340 nm, λ_EM_ 460 nm) and flavoprotein fluorescence (λ_EX_ 460 nm, λ_EM_ 520 nm), as shown in Figure 1D-1E.

### 2.4 Measuring *in situ* Mitochondrial ROS

Mitochondrial ROS generation was measured using the probe mitoSOX ^39, 40^ (tetraphenylphosphonium-conjugated dihydroethidium, TPP^+^–DHE) at a concentration of 1.5 µM, since the TPP^+^ moiety is known to uncouple mitochondria at concentrations ≥2.5 µM ^41-43^. Hearts subject to conditions (i-vii) described above were equilibrated for 10 min., loaded with 1.5 µM mitoSOX for 5 min. before immediately being subjected to 25 min. ischemia. Fluorescence was monitored at λ_EX_ 510 nm, λ_EM_ 580 nm.

Since the λ_EX_ and λ_EM_ of mitoSOX overlap with absorbance spectra of endogenous chromophores in cardiomyocytes (e.g., myoglobin and cytochromes), changes in chromophore absorbance during the course of IR could impact the fluorescent signal. Furthermore, the distribution of the mitoSOX probe between cytosolic and mitochondrial matrix compartments is determined by the mitochondrial membrane potential (ΔΨ_m_), which would also be expected to change during IR. To correct for these potential cofounding effects, a series of hearts were loaded with fully-oxidized mitoSOX ^37^, which fluoresces in a manner that is independent of ROS generation, but is still subject to the effects of chromophore absorbance and probe distribution. Fluorescent data from these hearts was used to correct data obtained with naïve mitoSOX (conditions i-vii), yielding a net signal that originated only from *in-situ* probe oxidation, without contribution from other factors. The data process for this correction is illustrated in Supplemental Figure 1, which also shows a minimal change in signal during IR when no mitoSOX was present. In addition, absorbance spectra of compounds tested in this study (AR, DMM, S1QEL, etc.) were also measured, to ensure these chemicals did not absorb significant amounts of light at the λ_EX_ and λ_EM_ of mitoSOX (Supplemental Figure 2).

### Measuring *in situ* Mitochondrial Membrane Potential (ΔΨ_m_)

In order to assess the timing of mitochondrial PT pore opening, mitochondrial membrane potential was measured using TMRE at 500 nM delivered for 20 min. prior to ischemia. The PT pore inhibitor CsA (0.8 µM) was optionally delivered for 5 min. prior to ischemia. To determine changes upon reperfusion, TMRE data were normalized to the fluorescent signal during the last 5 min. of ischemia. Two conditions were examined: control IR, and IR with MCT-1 inhibition (same as condition (vi) in section 2.2 above). Attempts to determine the effect of CsA on PT pore opening in the presence of AR-C155858 were confounded by interactions between these molecules and TMRE, resulting in precipitation of components in Krebs-Henseleit buffer.

### 2.6 Effluent Analysis by High-Performance Liquid Chromatography

Effluents from control and AR treated hearts were collected in 1 min. intervals for the first 3 min. of reperfusion, and immediately treated with 10 % perchloric acid. Following addition of 100 nmols butyrate as an internal standard, samples were frozen in liquid N_2_ and stored at -80°C until analysis. Samples were centrifuged at 20,000 x *g* to remove insoluble materials. Metabolites were resolved on HPLC (Shimadzu Prominence 20 system) using two 300 × 7.8 mm Aminex HPX-87H columns (BioRad, Carlsbad CA, USA) in series with 10 mM H_2_SO_4_ mobile phase (flow rate: 0.7 ml/min) and 100 µl sample injected on column. Succinate and lactate were detected using a photodiode array measuring absorbance at 210 nm as previously described ^2^. A standard curve was constructed for calibration. Lactate data were corrected for 1.2 mM lactate contained in the Krebs-Henseleit buffer.

### 2.7 Quantitation and Statistical Analysis

Comparisons between groups were made using ANOVA, followed by unpaired Student’s t-tests. Data are shown as means ± SEM. Numbers of biological replicates (N) are noted in the figures. Significance was set at α = 0.05.

## 3. RESULTS

### 3.1 Cardiac Fluorescence

Although similar spectrofluorometric cardiac perfusion apparatus has previously been constructed ^32, 44-48^, prior efforts have used larger animal hearts (rats, guinea pigs) or have not simultaneously measured cardiac function. To the best of our knowledge, this is the first study of mouse hearts with simultaneous fluorescence and functional assessment.

The spectrofluorimetric perfusion system was first validated by monitoring NAD(P)H and flavoprotein autofluorescence during ischemia and reperfusion (Fig. 1D and 1E). As expected, NAD(P)H (λ_EX_ 340 nm, λ_EM_ 460 nm) autofluorescence immediately rose upon ischemia, in agreement with previous reports ^32, 44, 49, 50^. Likewise, flavoprotein fluorescence (λ_Ex_ 460 nm, λ_Em_520 nm) decreased upon ischemia ^32, 50, 51^. Both parameters returned to baseline levels immediately upon reperfusion.

NAD(P)H autofluorescence in the heart is thought to mainly represent mitochondrial NADH, the substrate for Cx-I which accumulates during ischemia due to a highly reduced respiratory chain ^50^. The mitochondrial ATP synthase is thought to operate in reverse during ischemia, to maintain mitochondrial membrane potential (ΔΨ_m_) ^52, 53^, and this also contributes to feedback inhibition of Cx-I resulting in high [NADH] ^11^. Notably, a decrease in NAD(P)H signal was observed ∼12 min. into ischemia, concurrent with the onset of ischemic hyper-contracture (not shown). The latter is thought to indicate the onset of an energetic crisis (no ATP to relax contractile machinery) ^54^.

### 3.2 MitoSOX Fluorescence

Having validated the cardiac fluorescence system, we next sought to use it for measurement of ROS generation during IR. Figure 2A shows cardiac functional measurements throughout IR (heart rate x pressure product, RPP), while Figure 2B shows the corrected mitoSOX fluorescent readout over the same period.

**Figure 2.**
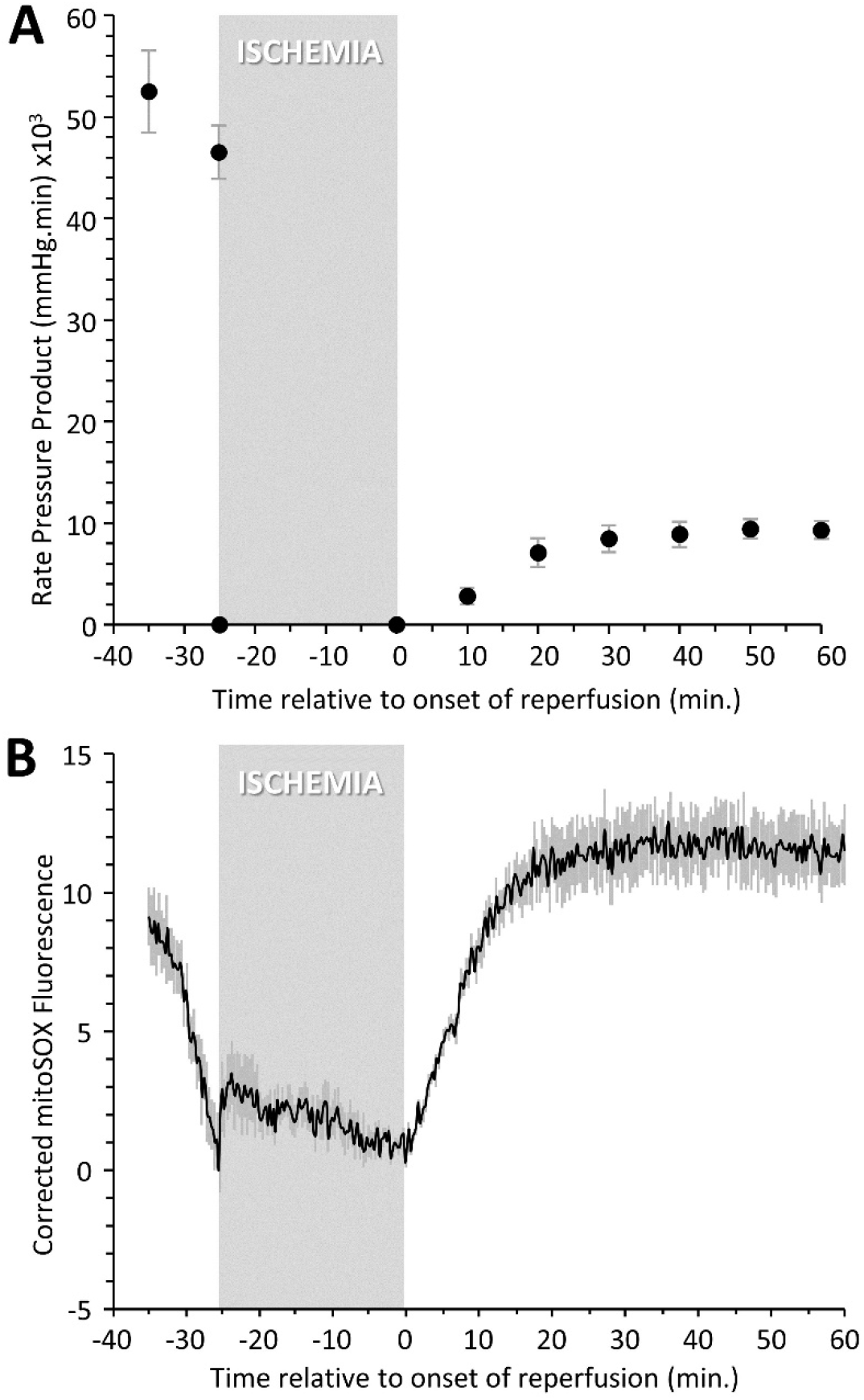
Cardiac Function vs. mitoSOX Fluorescence During IR. **(A):** Cardiac functional measurements. Graph shows heart rate x left ventricular developed pressure (i.e., rate pressure product, RPP). The period of 25 min. ischemia is indicated. Data are means ± SEM, N= 16. **(B):** Corrected mitoSOX fluorescence (510/580 nm) during IR. Fluorescent data were processed as described in the methods and as shown in Supplemental Figure 1, to correct for changes in absorbance of endogenous cytochromes at the fluorescence wavelengths. Data are means ± SEM, N= 6. <u>Note</u>: mitoSOX fluorescent measurements were not performed for the entire perfusion time for all hearts in panel A (e.g., some hearts were used for controls or other measurements), hence different N between panels.

To account for changes in the absorbance of endogenous chromophores during IR, or probe distribution, additional hearts were perfused with oxidized mitoSOX or with no probe at all. No significant change in the fluorescent signal was observed in hearts without mitoSOX (Supplemental Figure 1A). However, delivery of oxidized mitoSOX resulted in a rapid increase in the fluorescent signal during dye loading (Supplemental Figure 1B). At the onset of ischemia, an additional signal increase was seen. This is expected, since it has been shown that cardiac tissue absorbance at 510 nm and 580 nm decreases during ischemia ^55^. Upon reperfusion, after a small increase, the oxidized mitoSOX signal steadily declined for the remainder of the experiment. This is possibly due to loss of the dye from mitochondria as a result of changes in (ΔΨ_m_) as mitochondrial integrity becomes compromised.

Raw mitoSOX fluorescence traces (Supplemental Figure 1C) also increased slightly during loading, but not to the same extent as the fully oxidized probe. A sharp signal increase at the onset of ischemia may represent a burst of ROS generation as the terminal respiratory chain becomes inhibited. Upon reperfusion, a sustained signal increase was observed for ∼10 min., followed by a decline. Using the oxidized mitoSOX data to correct the raw mitoSOX data, Figure 2B (Supplemental Figure 1D) shows the signal resulting from oxidation of the probe during IR. Upon reperfusion, a sustained increase in the redox-dependent mitoSOX signal was seen, and this is consistent with the concept that a burst of mitochondrial ROS generation occurs during the first minutes of reperfusion ^1, 56-58^.

It has been posited that reverse electron transport (RET) at Cx-I is the primary source of ROS during reperfusion, and recently a novel series of inhibitors that target ROS generation at the ubiquinone (Q) binding site of Cx-I (termed S1QELs) were shown to elicit cardioprotection against IR injury ^59^. Succinate levels return to baseline within the first 5 min. of reperfusion ^2^, and accordingly, examining mitoSOX fluorescence during this time period revealed that the immediate signal increase at the onset of reperfusion in control hearts was suppressed in hearts treated with a S1QEL (Figure 3A-3C). These data suggest that the ROS signal detected by mitoSOX during the first minutes of reperfusion originates from Cx-I RET. However, by 2 minutes the rate of signal increase in S1QEL hearts had returned to that seen in control hearts, potentially indicating a role for other sources of ROS ^17^. While no effect of S1QEL on flavoprotein fluorescence was observed (Figure 3E), S1QEL did cause a slight detriment in the elevation of NAD(P)H fluorescence at the start of ischemia (37±4% with S1QEL vs. 53±5% in controls, p=0.042). However, it also blunted the NAD(P)H response to ischemic hyper-contracture (Figure 3D). The combination of these effects was such that the drop in NAD(P)H signal at the onset of reperfusion was not significantly different between S1QEL vs. control (30±4% vs. 36±4% respectively), suggesting that consumption of NADH by Cx-I during early reperfusion was not impacted by the compound.

**Figure 3.**
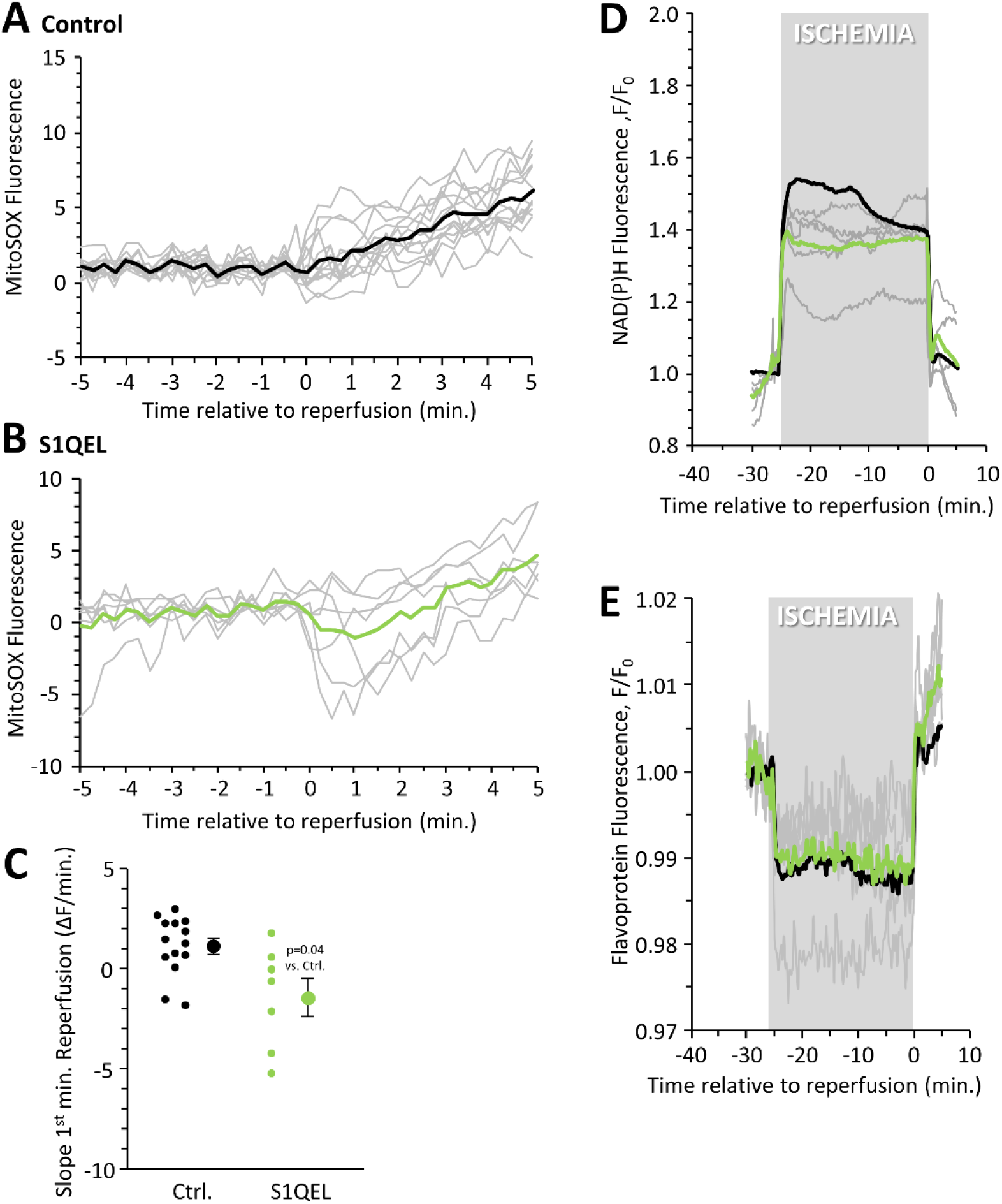
Impact of S1QEL on IR injury dynamics. Hearts were treated with the Cx-I Q-site ROS generation inhibitor S1QEL1.1 for 5 min. pre-and post-ischemia (Figure 1 schematic). **(A):** Control mitoSOX fluorescent data during the first 5 min. of reperfusion (from Figure 2B) for comparative purposes. **(B):** mitoSOX fluorescent data from S1QEL treated hearts. Gray traces show individual data, with averages shown as bold line. **(C):** Calculated slopes from 1st minute of reperfusion, for the data in panels A/B. For each condition, individual data points are shown on the left, with mean ± SEM on the right (N for each condition can be seen from number of data points). p values (ANOVA followed by unpaired Student’s t-test) for differences between groups are denoted. **(D):** NAD(P)H autofluorescence (340/460 nm) during ischemia and reperfusion. Traces in gray are individual hearts, with the average (N=5) shown in green. **(E):** Flavoprotein autofluorescence (462/520 nm) during ischemia and reperfusion. Traces in gray are individual hearts, with the average (N=5) shown in green. For panels D and E, the average data from control hearts (from Figure 1) is shown in black for comparative purposes. Fluorescence data are normalized to pre-ischemic levels.

### 3.3 Preventing Succinate Efflux Exacerbates Mitochondrial ROS at Reperfusion

Upon reperfusion of ischemic heart, ⅓ of accumulated succinate is oxidized by Cx-II, driving ROS generation ^2, 10^. Consistent with this, the Cx-II competitive inhibitor malonate is reported to be cardioprotective when delivered at reperfusion ^6, 7, 60^. The remaining ⅔ of succinate accumulated during ischemia is released from tissue upon reperfusion, and the pathway for this release was recently elucidated as monocarboxylate transporter 1 (MCT-1) ^2, 10^. To test the impact of MCT-1 inhibition on ROS generation at reperfusion, hearts were infused peri-ischemically with the MCT-1 inhibitor AR-C155858 (AR) ^61^, which resulted in a significant decrease in succinate release into the post-cardiac effluent during the first 3 minutes of reperfusion (Supplemental Figure 3). As expected, lactate efflux was also significantly diminished, confirming MCT-1 inhibition.

Figures 4C & 4F show that AR resulted in a significantly greater rate of ROS generation during the first minute of reperfusion, compared to control. To investigate the requirement for Cx-II in the additional AR-induced mitoSOX signal, the competitive Cx-II inhibitor dimethyl malonate (DMM, 5mM) was used. Surprisingly, DMM alone at this concentration did not significantly impact the mitoSOX signal, and it also did not significantly blunt the additional signal induced by AR (Figure 4B, 4D & 4F). We hypothesized that because malonate is a competitive Cx-II inhibitor, it may not be able to out-compete the additional succinate present in cells caused by MCT-1 inhibition. Supporting this hypothesis, tandem administration of a higher dose of DMM (10 mM) was capable of blocking the elevated mitoSOX signal elicited by AR, returning it to control levels (Figure 4E & 4F). Furthermore, the potent Cx-II inhibitor atpenin A5 (AA5, 100 nM) was also effective in blocking the additional mitoSOX signal induced by AR (Supplemental Figure 4A & 4B). Overall, these data suggest that MCT-1 inhibition enhances mitochondrial ROS generation in the first minute of reperfusion, in a manner that can be blocked by inhibitors of Cx-II.

**Figure 4.**
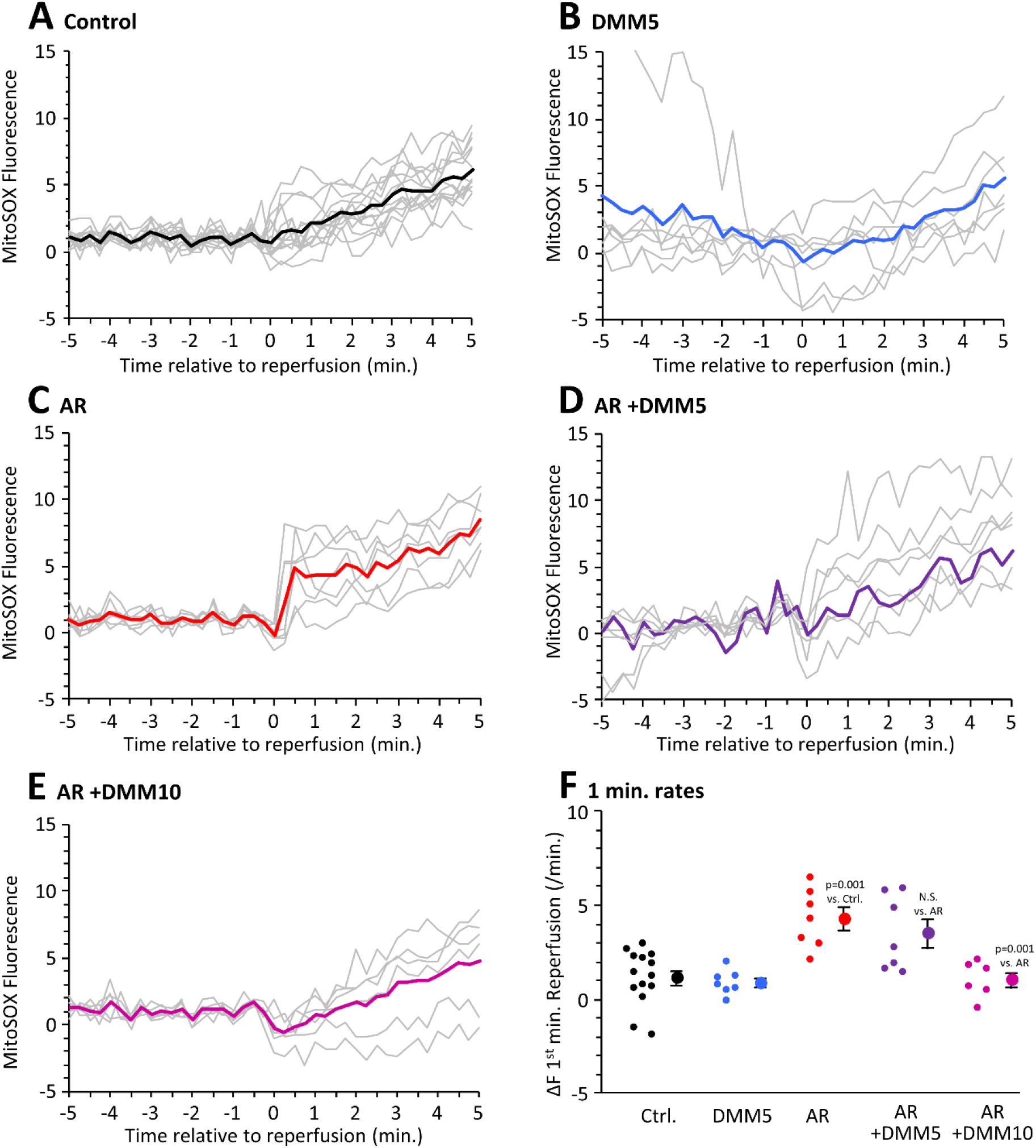
Impact of AR and Cx-II Inhibitors on mitoSOX Fluorescence at Reperfusion. **(A-E):** Corrected mitoSOX traces 5 min. pre-and post-reperfusion, for the labeled conditions. Gray traces are individual data, with averages shown as bold colored lines (see Figure 1 for color scheme). **(F):** Calculated slopes from 1^st^ minute of reperfusion, for the data in panels A-E. For each condition, individual data points are shown on the left, with mean ± SEM on the right (N for each condition can be seen from number of data points). p values (ANOVA followed by unpaired Student’s t-test) for differences between groups are denoted.

### 3.4 Blocking Succinate Release via MCT-1 Worsens IR Injury

In agreement with the role of succinate-derived ROS as a driver of post-IR pathology, DMM alone improved and AR worsened, IR injury (Figure 5A-C, cardiac functional recovery and infarct size). In addition, tandem administration of low dose DMM (5 mM) failed to reverse the impact of AR, whereas high dose DMM (10 mM) was protective. Furthermore, administration of the potent Cx-II inhibitor AA5 also blocked the impact of AR on functional recovery and infarct (Supplemental Figure 4C-E), and in-fact was more protective than AA5 alone ^2^. These effects of MCT-1 inhibition and Cx-II inhibition were more significant for myocardial infarction (Figure 5C) than for functional recovery (Figure 5B), but all trended in the same direction similar to mitoSOX data (Figure 4), i.e., interventions that increased mitoSOX signal at reperfusion worsened IR injury, and those that decreased it improved IR injury. Overall, a correlation was observed between the effect of interventions on mitoSOX at reperfusion and on infarct size and functional recovery, as illustrated in Figure 6.

**Figure 5.**
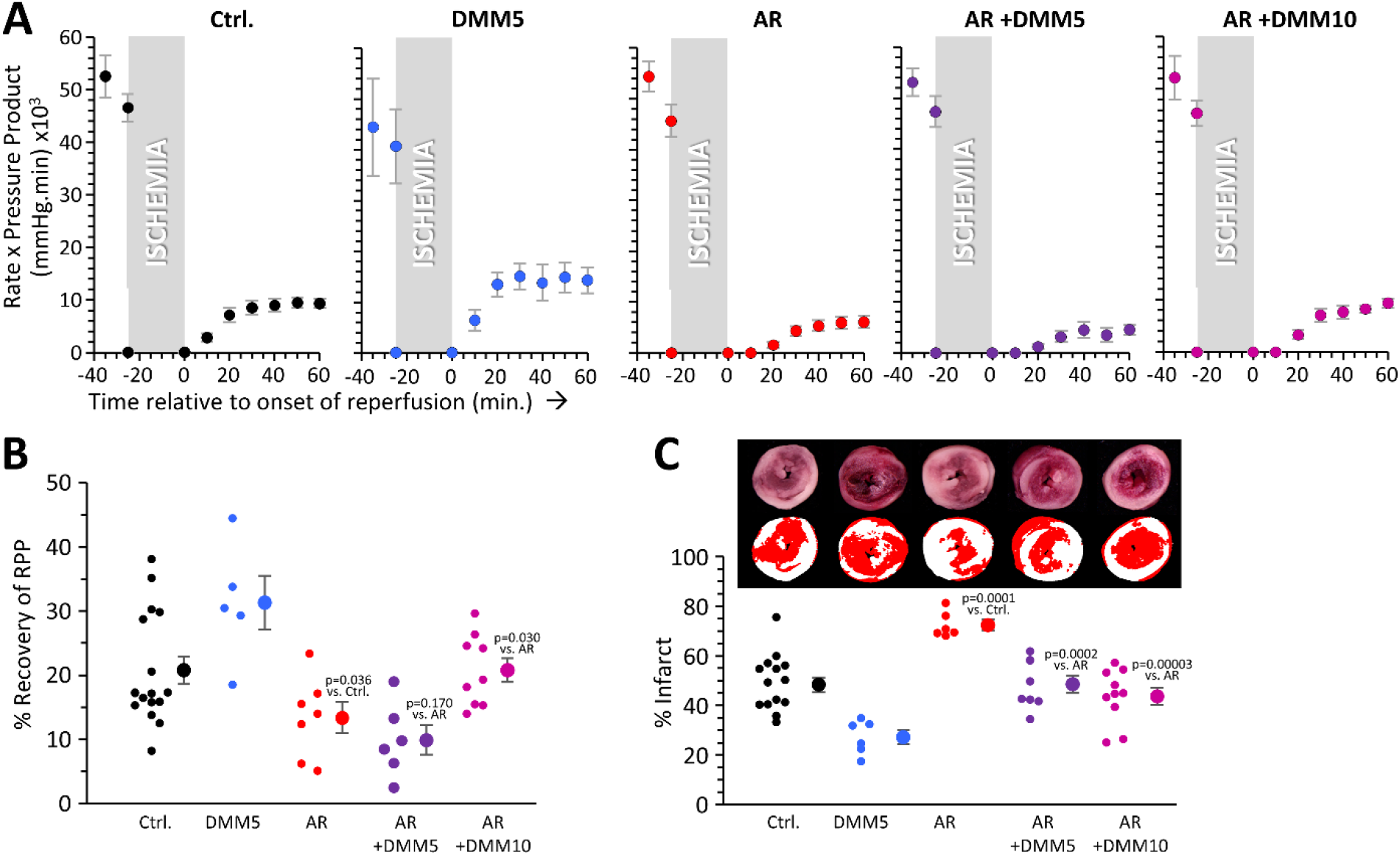
Impact of AR and Cx-II Inhibitors on Outcomes of IR Injury. **(A):** Cardiac functional measurements during IR. Graphs show heart rate x left ventricular developed pressure (i.e., rate pressure product, RPP) for each of the conditions examined. The period of 25 min. ischemia is indicated. Data are means ± SEM. **(B):** Quantitation of percent functional recovery, i.e., cardiac function at 60 min. of reperfusion as a percentage of that immediately before ischemia (−25 min. time point). For each condition, individual data points are shown on the left, with mean ± SEM shown on the right (N for each condition can be seen from number of data points). p values (ANOVA followed by unpaired Student’s t-test) for differences between groups are denoted above the data. **(C):** Myocardial infarct size for each condition. Images above the graph show representative TTC stained heart cross-sections, with pseudo-colored mask used for planimetry below. Graph shows data for each condition, with individual data points on the left, mean ± SEM on the right (N for each condition can be seen from number of data points). p values (ANOVA followed by unpaired Student’s t-test) for differences between groups are denoted above the data.

**Figure 6.**
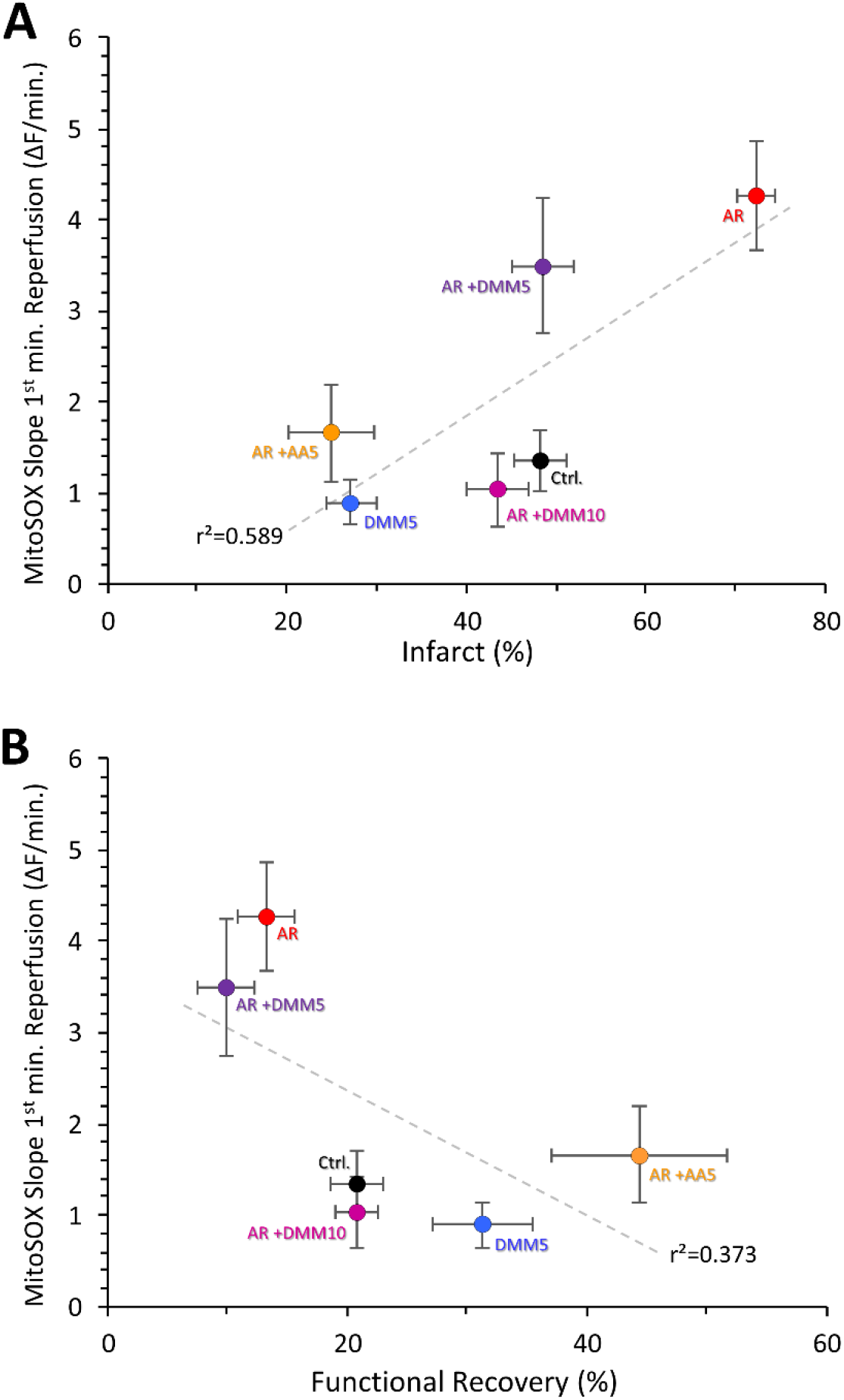
Correlations Between mitoSOX Fluorescence During Reperfusion and IR Outcomes. Graphs show correlation between mitoSOX slope data and **(A):** infarct size or **(B):** functional recovery. Each data point is the mean ± SEM for a condition (see color scheme in Figure 1). mitoSOX data are from Figure 3F and Supplemental Figure 5B. Infarct data are from Figure 4C and Supplemental Figure 5E. Functional data are from Figure 4B and Supplemental Figure 5D. Linear curve fits are shown, with correlation coefficient (r^2^) listed alongside.

### 3.5 Timing of Mitochondrial ROS at Reperfusion vs. PT Pore Opening

While it is largely accepted that ROS can trigger PT pore opening, it has also been proposed that during reperfusion injury, the PT pore itself may drive ROS generation ^23, 32^. To test this, we measured the effect of the PT pore inhibitor CsA on mitoSOX fluorescence during reperfusion. 0.8 µM CsA was chosen, as this was previously shown to elicit protection in mouse hearts ^38^. As shown in Figure 7B & 7C, CsA had no impact on mitoSOX fluorescence during reperfusion. To confirm that PT pore opening did occur, hearts were loaded with the ΔΨ_m_ indicator TMRE. Upon reperfusion, an immediate rise in TMRE fluorescence was observed. Subsequently, in control hearts the TMRE signal declined from ∼5 min. into reperfusion, whereas in CsA treated hearts the signal was sustained (Figure 7D & 7E). We thus infer that PT pore opening occurs no sooner than 5 min. into reperfusion. Despite a small blip in the mitoSOX signal in control hearts at ∼6.5 min., no substantial difference was seen between control & CsA hearts at this time point, concurrent with divergence of the TMRE traces, thus suggesting no secondary ROS burst due to PT pore opening.

**Figure 7.**
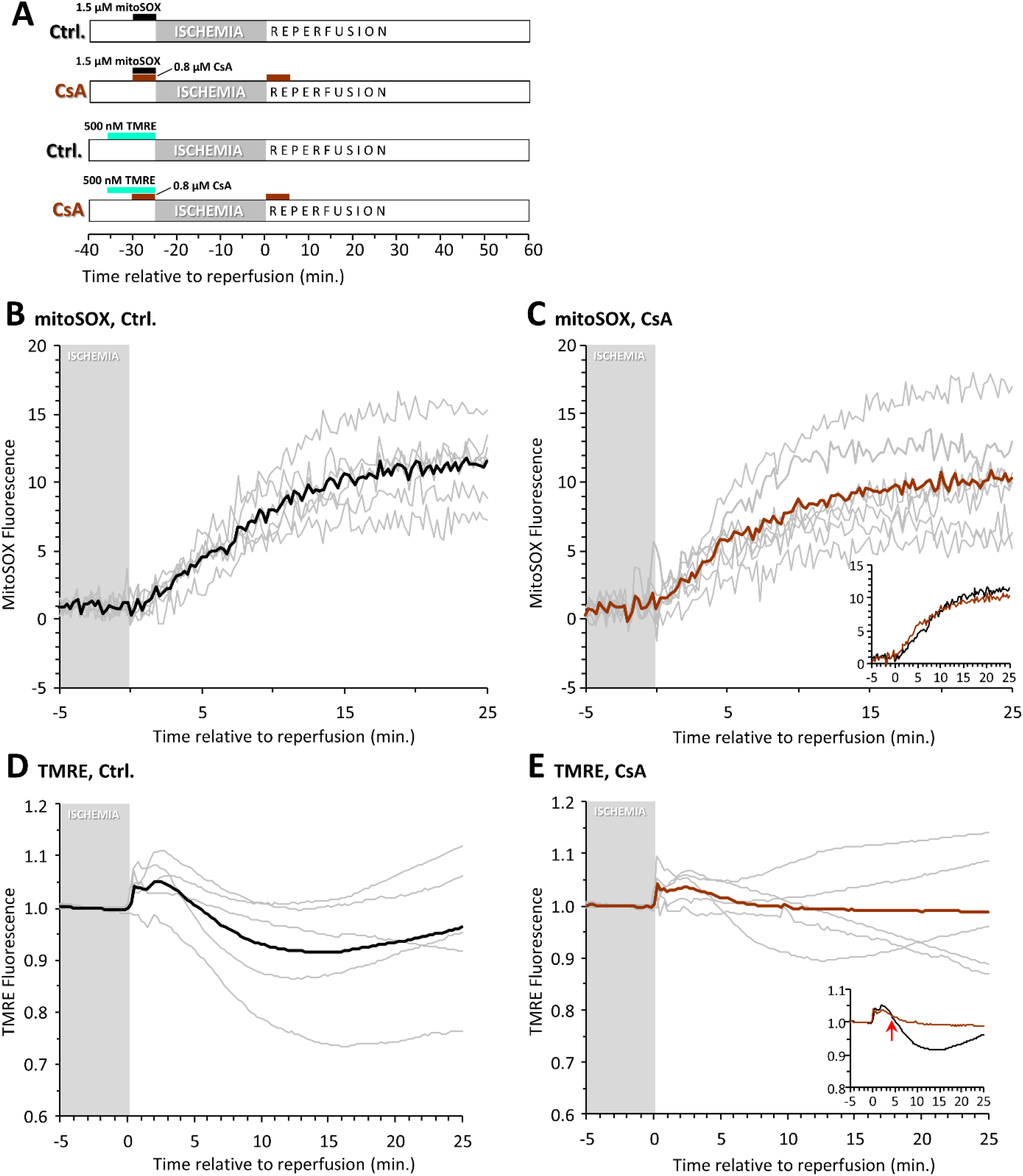
Temporal Relationship Between mitoSOX and PT Pore Opening During IR. **(A):** Schematic showing perfusion conditions. Where indicated, mitoSOX (ROS indicator), TMRE (ΔΨ_m_ indicator) and/or CsA (PT pore inhibitor) were administered at the listed concentrations. **(B/C):** mitoSOX signal during early reperfusion under control or CsA condition. The last 5 min. of the ischemic period is indicated. Gray traces show individual data, with averages shown as bold lines. Inset to panel C shows averages for control and CsA superimposed. **(D/E):** Normalized TMRE fluorescence during early reperfusion under control or CsA condition. The last 5 min. of the ischemic period is indicated. Gray traces show individual data, with averages shown as bold lines. Inset to panel C shows averages for control and CsA superimposed. Red arrow indicates the point at which traces diverge, 5 min. into reperfusion.

Finally, as expected from its impact on mitoSOX fluorescence during early reperfusion (Figure 4), inhibition of MCT-1 led to an accelerated loss of the TMRE signal during reperfusion, indicating faster opening of the PT pore (Supplemental Figure 5).

## 4. DISCUSSION

Cardiac IR is a complex pathology, with numerous links between early and late events in the development of myocardial infarction. Shortly after reperfusion, a burst of ROS generation and mitochondrial Ca^2+^ overload both trigger opening of the mitochondrial PT pore, a key event in necrotic cell death of cardiomyocytes. Following this, inflammatory cells (macrophages, neutrophils) are recruited to the heart, where they mediate responses that lead to cardiac remodeling, fibrosis, and eventual development of hypertrophy and heart failure ^62-64^.

Accumulation of succinate in hypoxia/ischemia is highly conserved ^3, 4^, but this succinate drives ROS generation upon tissue reperfusion, and this has led to a consensus that <u>intra</u>cellular succinate plays a detrimental role in reperfusion injury ^1, 2, 16, 65, 66^. In contrast, succinate release from cells via MCT-1 ^2, 10, 23^, along with the recent identification of a succinate receptor (GPR91/SUCNR1 ^25^), have led to the notion that <u>extra</u>cellular succinate may also play a role in IR ^6, 10^. In this regard, findings from an *in-vivo* model of cardiac IR injury demonstrated that blocking succinate release via MCT-1 was cardioprotective, likely due to inhibiting immune system activation ^6, 10^. However, herein our data reveal that acutely blocking MCT-1 in a perfused heart system (where there are no inflammatory cells) leads to greater ROS generation and the worsening of IR injury. Reconciling these findings, it is possible that MCT-1 inhibition is indeed detrimental in the acute setting at the level of cardiomyocytes, but this is balanced *in-vivo* by a longer-term effect of MCT-1 inhibition, possibly involving receptor-mediated succinate effects on other cell types, including inflammatory cells. Together, these studies highlight that the therapeutic targeting of MCT-1 may require careful timing and titration, to balance detrimental vs. beneficial effects of blocking succinate release.

Inhibiting MCT-1 in the heart lowered succinate release into the effluent by ∼30%. Assuming this succinate was retained in cells and available for oxidation by Cx-II, it is therefore not surprising that MCT-1 inhibition also led to enhanced ROS generation. Furthermore, while 5 mM of the competitive Cx-II inhibitor malonate was ineffective at blocking this additional ROS, doubling its concentration to 10 mM effectively overrode the impact of AR. Unfortunately, due to the physical nature of dimethylmalonate (liquid) and our drug infusion system, it was not possible to test additional even higher doses of DMM. Instead, we tested the potent Cx-II inhibitor atpenin A5 (AA5, IC_50_ ∼10 nM ^67^), and found that it was able to completely abrogate the additional ROS induced by AR, and this resulted in cardioprotection to a level greater than baseline control IR injury (Supplementary Figure 4). In-fact, the combination of AR and AA5 may be considered optimal from a therapeutic perspective – keeping succinate inside cells to prevent its signaling effects, and preventing its oxidation to generate ROS.

Although there is a general consensus that mitochondrial ROS generation during early reperfusion is an upstream event that triggers PT pore opening, it has also been suggested that pore opening itself is the main driver of ROS generation in early reperfusion ^23, 32^. Our results (Figure 7) suggest that the burst of ROS at reperfusion is independent of the PT pore, since evidence for pore opening (i.e., a CsA-sensitive loss of ΔΨ_m_) was not observed until at least 5 min. into reperfusion, by which time a large amount of ROS generation had already occurred.

It should not go unmentioned that Ca^2+^ is also considered to be a major trigger for PT pore opening ^68, 69^, with both Ca^2+^ and ROS thought to potentiate each other’s effects at promoting pore formation ^70^. Thus, future experiments should be directed at using fluorescent Ca^2+^ probes in this or similar perfusion systems, to understand *in-situ* Ca^2+^ kinetics and how they relate to ROS and pore opening kinetics in early reperfusion.

A number of caveats regarding the use of mitoSOX as a probe for mitochondrial ROS should be addressed. Firstly, it is known the probe can be oxidized by reactants other than ROS, although notably such one-electron oxidations result in non-fluorescent products that would be undetectable in our measurement system ^71^. Furthermore, the mitoSOX signal increase in early reperfusion was inhibited by S1QEL, a specific inhibitor of ROS generation at Cx-I ^59^, so we consider any contribution from other poorly-characterized oxidation sources to be minimal. Secondly, the mitoSOX signal is impacted both by its distribution between extracellular, cytosolic and mitochondrial compartments ^37^, and by the primary and secondary filter effects from endogenous chromophores ^47, 55^ (see methods section 2.4). However, our use of fully-oxidized mitoSOX as a control (Supplementary Figure 1) ensures that any changes in the mitoSOX signal we observed originated only from *in-situ* probe oxidation, and not from redistribution of the probe or changes in the absorbance of myoglobin, cytochromes, etc.

While mitoSOX does confer a number of limitations regarding the assignment of the signal to a particular reactive oxygen species, similar issues of probe specificity are also broadly applicable to genetically encoded biosensors ^72, 73^. In addition, more precise analytical methods such as LC-MS separation of mitoSOX oxidation products ^37, 74^ require time-consuming isolation steps that may release components that further oxidize the probe during isolation, and so may not be readily compatible with the rapid kinetics of the events observed herein. The development of more precise genetically-encoded and non ΔΨ_m_ dependent mitochondrial ROS probes may therefore be useful in future studies.

Overall, the findings herein demonstrate that blocking MCT-1 in cardiac IR leads to inhibition of succinate release, which worsens IR injury due to enhanced mitochondrial ROS generation. Concurrent inhibition of Cx-II abrogates these effects, highlighting the importance of succinate oxidation at Cx-II in the pathology of IR injury. Furthermore, these events appear to lie temporally upstream of PT pore opening. Future experiments could explore the role of local succinate signaling in the heart, to determine if SUCNR1 may modulate responses to IR in a manner independent of inflammatory cells, providing further insight on the delicate balance of succinate dynamics in IR injury.

## Author Contribution Statement

ASM and PSB designed the research. ASM and SMN performed experiments. ASM and PSB analyzed the data and wrote the manuscript. All authors approved the final version of the manuscript.

## Acknowledgements

We thank Robert S. Balaban (NHLBI) for constructive discussions regarding corrections for primary and secondary filtering effects, during early design stages of this project.

## Data Availability Statement

All original data used to prepare the figures is available in spreadsheet form, on the data sharing website FigShare (https://doi.org/10.6084/m9.figshare.19319627, to be unembargoed upon publication).

## REFERENCES

1. Chouchani ET, Pell VR, Gaude E, Aksentijevic D, Sundier SY, Robb EL, Logan A, Nadtochiy SM, Ord ENJ, Smith AC, Eyassu F, Shirley R, Hu CH, Dare AJ, James AM, Rogatti S, Hartley RC, Eaton S, Costa ASH, Brookes PS, Davidson SM, Duchen MR, Saeb-Parsy K, Shattock MJ, Robinson AJ, Work LM, Frezza C, Krieg T, Murphy MP. Ischaemic accumulation of succinate controls reperfusion injury through mitochondrial ros. Nature. 2014;515:431–435

2. Zhang J, Wang YT, Miller JH, Day MM, Munger JC, Brookes PS. Accumulation of succinate in cardiac ischemia primarily occurs via canonical krebs cycle activity. Cell Rep. 2018;23:2617–2628

3. Hochachka PW, Dressendorfer RH. Succinate accumulation in man during exercise. Eur J Appl Physiol Occup Physiol. 1976;35:235–242

4. Hochachka PW, Owen TG, Allen JF, Whittow GC. Multiple end products of anaerobiosis in diving vertebrates. Comp Biochem Physiol B. 1975;50:17–22

5. Pell VR, Chouchani ET, Frezza C, Murphy MP, Krieg T. Succinate metabolism: A new therapeutic target for myocardial reperfusion injury. Cardiovasc Res. 2016;111:134–141

6. Kula-Alwar D, Prag HA, Krieg T. Targeting succinate metabolism in ischemia/reperfusion injury. Circulation. 2019;140:1968–1970

7. Valls-Lacalle L, Barba I, Miro-Casas E, Ruiz-Meana M, Rodriguez-Sinovas A, Garcia-Dorado D. Selective inhibition of succinate dehydrogenase in reperfused myocardium with intracoronary malonate reduces infarct size. Sci Rep. 2018;8:2442

8. Pell VR, Chouchani ET, Murphy MP, Brookes PS, Krieg T. Moving forwards by blocking back-flow: The yin and yang of mi therapy. Circ Res. 2016;118:898–906

9. Chouchani ET, Pell VR, James AM, Work LM, Saeb-Parsy K, Frezza C, Krieg T, Murphy MP. A unifying mechanism for mitochondrial superoxide production during ischemia-reperfusion injury. Cell Metab. 2016;23:254–263

10. Prag HA, Gruszczyk AV, Huang MM, Beach TE, Young T, Tronci L, Nikitopoulou E, Mulvey JF, Ascione R, Hadjihambi A, Shattock MJ, Pellerin L, Saeb-Parsy K, Frezza C, James AM, Krieg T, Murphy MP, Aksentijevic D. Mechanism of succinate efflux upon reperfusion of the ischaemic heart. Cardiovasc Res. 2021;117:1188–1201

11. Lambert AJ, Brand MD. Superoxide production by nadh:Ubiquinone oxidoreductase (complex i) depends on the ph gradient across the mitochondrial inner membrane. Biochem J. 2004;382:511–517

12. Watson MA, Wong HS, Brand MD. Use of s1qels and s3qels to link mitochondrial sites of superoxide and hydrogen peroxide generation to physiological and pathological outcomes. Biochem Soc Trans. 2019;47:1461–1469

13. Wong HS, Monternier PA, Brand MD. S1qels suppress mitochondrial superoxide/hydrogen peroxide production from site iq without inhibiting reverse electron flow through complex i. Free Radic Biol Med. 2019;143:545–559

14. Cadenas E, Boveris A, Ragan CI, Stoppani AO. Production of superoxide radicals and hydrogen peroxide by nadh-ubiquinone reductase and ubiquinol-cytochrome c reductase from beef-heart mitochondria. Arch Biochem Biophys. 1977;180:248–257

15. Turrens JF, Alexandre A, Lehninger AL. Ubisemiquinone is the electron donor for superoxide formation by complex iii of heart mitochondria. Arch Biochem Biophys. 1985;237:408–414

16. Murphy MP. How mitochondria produce reactive oxygen species. Biochem J. 2009;417:1–13

17. Chen Q, Vazquez EJ, Moghaddas S, Hoppel CL, Lesnefsky EJ. Production of reactive oxygen species by mitochondria: Central role of complex iii. J Biol Chem. 2003;278:36027–36031

18. Chen Q, Moghaddas S, Hoppel CL, Lesnefsky EJ. Ischemic defects in the electron transport chain increase the production of reactive oxygen species from isolated rat heart mitochondria. Am J Physiol Cell Physiol. 2008;294:C460–466

19. Morciano G, Giorgi C, Bonora M, Punzetti S, Pavasini R, Wieckowski MR, Campo G, Pinton P. Molecular identity of the mitochondrial permeability transition pore and its role in ischemia-reperfusion injury. J Mol Cell Cardiol. 2015;78:142–153

20. Bernardi P, Di Lisa F. The mitochondrial permeability transition pore: Molecular nature and role as a target in cardioprotection. J Mol Cell Cardiol. 2015;78:100–106

21. Beutner G, Alavian KN, Jonas EA, Porter GA, Jr. The mitochondrial permeability transition pore and atp synthase. Handb Exp Pharmacol. 2017;240:21–46

22. Halestrap AP, Richardson AP. The mitochondrial permeability transition: A current perspective on its identity and role in ischaemia/reperfusion injury. J Mol Cell Cardiol. 2015;78:129–141

23. Andrienko TN, Pasdois P, Pereira GC, Ovens MJ, Halestrap AP. The role of succinate and ros in reperfusion injury - a critical appraisal. J Mol Cell Cardiol. 2017;110:1–14

24. Reddy A, Bozi LHM, Yaghi OK, Mills EL, Xiao H, Nicholson HE, Paschini M, Paulo JA, Garrity R, Laznik-Bogoslavski D, Ferreira JCB, Carl CS, Sjoberg KA, Wojtaszewski JFP, Jeppesen JF, Kiens B, Gygi SP, Richter EA, Mathis D, Chouchani ET. Ph-gated succinate secretion regulates muscle remodeling in response to exercise. Cell. 2020;183:62–75 e17

25. He W, Miao FJ, Lin DC, Schwandner RT, Wang Z, Gao J, Chen JL, Tian H, Ling L. Citric acid cycle intermediates as ligands for orphan g-protein-coupled receptors. Nature. 2004;429:188–193

26. Tannahill GM, Curtis AM, Adamik J, Palsson-McDermott EM, McGettrick AF, Goel G, Frezza C, Bernard NJ, Kelly B, Foley NH, Zheng L, Gardet A, Tong Z, Jany SS, Corr SC, Haneklaus M, Caffrey BE, Pierce K, Walmsley S, Beasley FC, Cummins E, Nizet V, Whyte M, Taylor CT, Lin H, Masters SL, Gottlieb E, Kelly VP, Clish C, Auron PE, Xavier RJ, O’Neill LA. Succinate is an inflammatory signal that induces il-1beta through hif-1alpha. Nature. 2013;496:238–242

27. Trauelsen M, Hiron TK, Lin D, Petersen JE, Breton B, Husted AS, Hjorth SA, Inoue A, Frimurer TM, Bouvier M, O’Callaghan CA, Schwartz TW. Extracellular succinate hyperpolarizes m2 macrophages through sucnr1/gpr91-mediated gq signaling. Cell Rep. 2021;35:109246

28. Wu JY, Huang TW, Hsieh YT, Wang YF, Yen CC, Lee GL, Yeh CC, Peng YJ, Kuo YY, Wen HT, Lin HC, Hsiao CW, Wu KK, Kung HJ, Hsu YJ, Kuo CC. Cancer-derived succinate promotes macrophage polarization and cancer metastasis via succinate receptor. Mol Cell. 2020;77:213–227 e215

29. van Diepen JA, Robben JH, Hooiveld GJ, Carmone C, Alsady M, Boutens L, Bekkenkamp-Grovenstein M, Hijmans A, Engelke UFH, Wevers RA, Netea MG, Tack CJ, Stienstra R, Deen PMT. Sucnr1-mediated chemotaxis of macrophages aggravates obesity-induced inflammation and diabetes. Diabetologia. 2017;60:1304–1313

30. Rubic T, Lametschwandtner G, Jost S, Hinteregger S, Kund J, Carballido-Perrig N, Schwarzler C, Junt T, Voshol H, Meingassner JG, Mao X, Werner G, Rot A, Carballido JM. Triggering the succinate receptor gpr91 on dendritic cells enhances immunity. Nat Immunol. 2008;9:1261–1269

31. Vujic A, Koo ANM, Prag HA, Krieg T. Mitochondrial redox and tca cycle metabolite signaling in the heart. Free Radic Biol Med. 2021;166:287–296

32. Andrienko T, Pasdois P, Rossbach A, Halestrap AP. Real-time fluorescence measurements of ros and [ca2+] in ischemic / reperfused rat hearts: Detectable increases occur only after mitochondrial pore opening and are attenuated by ischemic preconditioning. PLoS One. 2016;11:e0167300

33. Cope DK, Impastato WK, Cohen MV, Downey JM. Volatile anesthetics protect the ischemic rabbit myocardium from infarction. Anesthesiology. 1997;86:699–709

34. Headrick JP, See Hoe LE, Du Toit EF, Peart JN. Opioid receptors and cardioprotection - ‘opioidergic conditioning’ of the heart. Br J Pharmacol. 2015;172:2026–2050

35. Molojavyi A, Preckel B, Comfere T, Mullenheim J, Thamer V, Schlack W. Effects of ketamine and its isomers on ischemic preconditioning in the isolated rat heart. Anesthesiology. 2001;94:623-629; discussion 625A-626A

36. Kishikawa JI, Inoue Y, Fujikawa M, Nishimura K, Nakanishi A, Tanabe T, Imamura H, Yokoyama K. General anesthetics cause mitochondrial dysfunction and reduction of intracellular atp levels. PLoS One. 2018;13:e0190213

37. Zielonka J, Vasquez-Vivar J, Kalyanaraman B. Detection of 2-hydroxyethidium in cellular systems: A unique marker product of superoxide and hydroethidine. Nat Protoc. 2008;3:8–21

38. Li J, Iorga A, Sharma S, Youn JY, Partow-Navid R, Umar S, Cai H, Rahman S, Eghbali M. Intralipid, a clinically safe compound, protects the heart against ischemia-reperfusion injury more efficiently than cyclosporine-a. Anesthesiology. 2012;117:836–846

39. Robinson KM, Janes MS, Pehar M, Monette JS, Ross MF, Hagen TM, Murphy MP, Beckman JS. Selective fluorescent imaging of superoxide in vivo using ethidium-based probes. Proc Natl Acad Sci U S A. 2006;103:15038–15043

40. Polster BM, Nicholls DG, Ge SX, Roelofs BA. Use of potentiometric fluorophores in the measurement of mitochondrial reactive oxygen species. Methods Enzymol. 2014;547:225–250

41. Roelofs BA, Ge SX, Studlack PE, Polster BM. Low micromolar concentrations of the superoxide probe mitosox uncouple neural mitochondria and inhibit complex iv. Free Radic Biol Med. 2015;86:250–258

42. Kulkarni CA, Fink BD, Gibbs BE, Chheda PR, Wu M, Sivitz WI, Kerns RJ. A novel triphenylphosphonium carrier to target mitochondria without uncoupling oxidative phosphorylation. J Med Chem. 2021;64:662–676

43. Reily C, Mitchell T, Chacko BK, Benavides G, Murphy MP, Darley-Usmar V. Mitochondrially targeted compounds and their impact on cellular bioenergetics. Redox Biol. 2013;1:86–93

44. Varadarajan SG, An J, Novalija E, Smart SC, Stowe DF. Changes in [na(+)](i), compartmental [ca(2+)], and nadh with dysfunction after global ischemia in intact hearts. Am J Physiol Heart Circ Physiol. 2001;280:H280–293

45. Riess ML, Camara AK, Chen Q, Novalija E, Rhodes SS, Stowe DF. Altered nadh and improved function by anesthetic and ischemic preconditioning in guinea pig intact hearts. Am J Physiol Heart Circ Physiol. 2002;283:H53–60

46. Lu LS, Liu YB, Sun CW, Lin LC, Su MJ, Wu CC. Optical mapping of myocardial reactive oxygen species production throughout the reperfusion of global ischemia. J Biomed Opt. 2006;11:021012

47. Bauer TM, Giles AV, Sun J, Femnou A, Covian R, Murphy E, Balaban RS. Perfused murine heart optical transmission spectroscopy using optical catheter and integrating sphere: Effects of ischemia/reperfusion. Anal Biochem. 2019;586:113443

48. Girouard SD, Pastore JM, Laurita KR, Gregory KW, Rosenbaum DS. Optical mapping in a new guinea pig model of ventricular tachycardia reveals mechanisms for multiple wavelengths in a single reentrant circuit. Circulation. 1996;93:603–613

49. Wengrowski AM, Kuzmiak-Glancy S, Jaimes R, 3rd, Kay MW. Nadh changes during hypoxia, ischemia, and increased work differ between isolated heart preparations. Am J Physiol Heart Circ Physiol. 2014;306:H529–537

50. Barlow CH, Harden WR, 3rd, Harken AH, Simson MB, Haselgrove JC, Chance B, O’Connor M, Austin G. Fluorescence mapping of mitochondrial redox changes in heart and brain. Crit Care Med. 1979;7:402–406

51. Chance B, Cohen P, Jobsis F, Schoener B. Intracellular oxidation-reduction states in vivo. Science. 1962;137:499–508

52. Ivanes F, Faccenda D, Gatliff J, Ahmed AA, Cocco S, Cheng CH, Allan E, Russell C, Duchen MR, Campanella M. The compound btb06584 is an if1 -dependent selective inhibitor of the mitochondrial f1 fo-atpase. Br J Pharmacol. 2014;171:4193–4206

53. Faccenda D, Campanella M. Molecular regulation of the mitochondrial f(1)f(o)-atpsynthase: Physiological and pathological significance of the inhibitory factor 1 (if(1)). Int J Cell Biol. 2012;2012:367934

54. Honda HM, Korge P, Weiss JN. Mitochondria and ischemia/reperfusion injury. Ann N Y Acad Sci. 2005;1047:248–258

55. Kanaide H, Yoshimura R, Makino N, Nakamura M. Regional myocardial function and metabolism during acute coronary artery occlusion. Am J Physiol. 1982;242:H980–989

56. Guarnieri C, Flamigni F, Caldarera CM. Role of oxygen in the cellular damage induced by re-oxygenation of hypoxic heart. J Mol Cell Cardiol. 1980;12:797–808

57. Hess ML, Manson NH. Molecular oxygen: Friend and foe. The role of the oxygen free radical system in the calcium paradox, the oxygen paradox and ischemia/reperfusion injury. J Mol Cell Cardiol. 1984;16:969–985

58. Raedschelders K, Ansley DM, Chen DD. The cellular and molecular origin of reactive oxygen species generation during myocardial ischemia and reperfusion. Pharmacol Ther. 2012;133:230–255

59. Brand MD, Goncalves RL, Orr AL, Vargas L, Gerencser AA, Borch Jensen M, Wang YT, Melov S, Turk CN, Matzen JT, Dardov VJ, Petrassi HM, Meeusen SL, Perevoshchikova IV, Jasper H, Brookes PS, Ainscow EK. Suppressors of superoxide-h2o2 production at site iq of mitochondrial complex i protect against stem cell hyperplasia and ischemia-reperfusion injury. Cell Metab. 2016;24:582–592

60. Valls-Lacalle L, Barba I, Miro-Casas E, Alburquerque-Bejar JJ, Ruiz-Meana M, Fuertes-Agudo M, Rodriguez-Sinovas A, Garcia-Dorado D. Succinate dehydrogenase inhibition with malonate during reperfusion reduces infarct size by preventing mitochondrial permeability transition. Cardiovasc Res. 2016;109:374–384

61. Ovens MJ, Davies AJ, Wilson MC, Murray CM, Halestrap AP. Ar-c155858 is a potent inhibitor of monocarboxylate transporters mct1 and mct2 that binds to an intracellular site involving transmembrane helices 7-10. Biochem J. 2010;425:523–530

62. Zuidema MY, Zhang C. Ischemia/reperfusion injury: The role of immune cells. World J Cardiol. 2010;2:325–332

63. Smiley D, Smith MA, Carreira V, Jiang M, Koch SE, Kelley M, Rubinstein J, Jones WK, Tranter M. Increased fibrosis and progression to heart failure in mrl mice following ischemia/reperfusion injury. Cardiovasc Pathol. 2014;23:327–334

64. Bonaventura A, Montecucco F, Dallegri F. Cellular recruitment in myocardial ischaemia/reperfusion injury. Eur J Clin Invest. 2016;46:590–601

65. Chen YR, Zweier JL. Cardiac mitochondria and reactive oxygen species generation. Circ Res. 2014;114:524–537

66. Quinlan CL, Perevoshchikova IV, Hey-Mogensen M, Orr AL, Brand MD. Sites of reactive oxygen species generation by mitochondria oxidizing different substrates. Redox Biol. 2013;1:304–312

67. Miyadera H, Shiomi K, Ui H, Yamaguchi Y, Masuma R, Tomoda H, Miyoshi H, Osanai A, Kita K, Omura S. Atpenins, potent and specific inhibitors of mitochondrial complex ii (succinate-ubiquinone oxidoreductase). Proc Natl Acad Sci U S A. 2003;100:473–477

68. Miyamae M, Camacho SA, Weiner MW, Figueredo VM. Attenuation of postischemic reperfusion injury is related to prevention of [ca2+]m overload in rat hearts. Am J Physiol. 1996;271:H2145–2153

69. Silverman HS, Stern MD. Ionic basis of ischaemic cardiac injury: Insights from cellular studies. Cardiovasc Res. 1994;28:581–597

70. Brookes PS, Yoon Y, Robotham JL, Anders MW, Sheu SS. Calcium, atp, and ros: A mitochondrial love-hate triangle. Am J Physiol Cell Physiol. 2004;287:C817–833

71. Zielonka J, Srinivasan S, Hardy M, Ouari O, Lopez M, Vasquez-Vivar J, Avadhani NG, Kalyanaraman B. Cytochrome c-mediated oxidation of hydroethidine and mito-hydroethidine in mitochondria: Identification of homo-and heterodimers. Free Radic Biol Med. 2008;44:835–846

72. Pak VV, Ezerina D, Lyublinskaya OG, Pedre B, Tyurin-Kuzmin PA, Mishina NM, Thauvin M, Young D, Wahni K, Martinez Gache SA, Demidovich AD, Ermakova YG, Maslova YD, Shokhina AG, Eroglu E, Bilan DS, Bogeski I, Michel T, Vriz S, Messens J, Belousov VV. Ultrasensitive genetically encoded indicator for hydrogen peroxide identifies roles for the oxidant in cell migration and mitochondrial function. Cell Metab. 2020;31:642–653 e646

73. Zhao Y, Hu Q, Cheng F, Su N, Wang A, Zou Y, Hu H, Chen X, Zhou HM, Huang X, Yang K, Zhu Q, Wang X, Yi J, Zhu L, Qian X, Chen L, Tang Y, Loscalzo J, Yang Y. Sonar, a highly responsive nad+/nadh sensor, allows high-throughput metabolic screening of anti-tumor agents. Cell Metab. 2015;21:777–789

74. Kalinovic S, Oelze M, Kroller-Schon S, Steven S, Vujacic-Mirski K, Kvandova M, Schmal I, Al Zuabi A, Munzel T, Daiber A. Comparison of mitochondrial superoxide detection ex vivo/in vivo by mitosox hplc method with classical assays in three different animal models of oxidative stress. Antioxidants (Basel). 2019;8

