## Supplemental Figs 1-5 for "Inhibiting succinate release worsens cardiac reperfusion injury by enhancing mitochondrial reactive oxygen species generation"

### SUPPLEMENTAL INFORMATION

#### 5 Figures & Legends

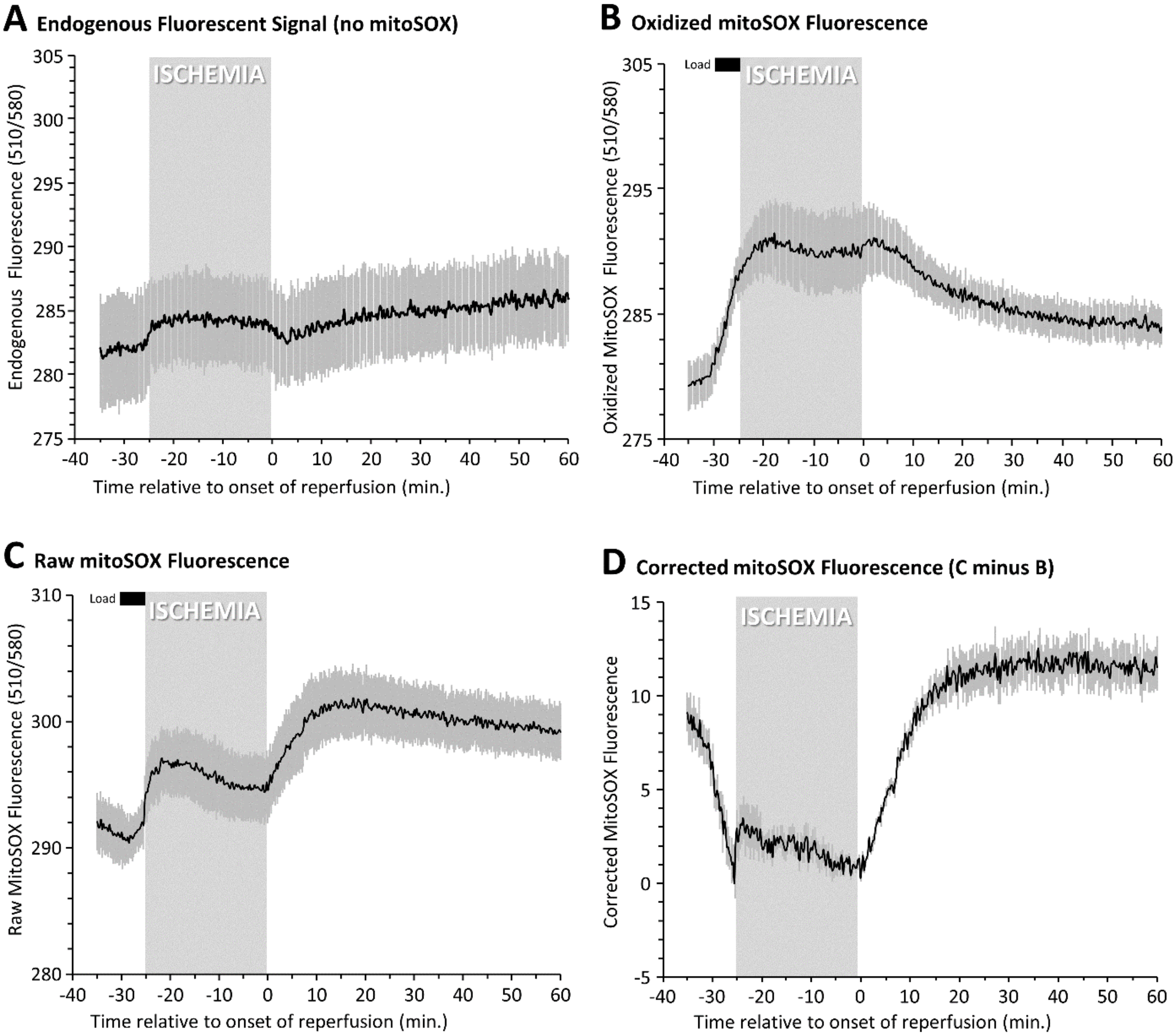


**Supplemental Figure 1. Correction of mitoSOX Fluorescence Data.** Fluorescent data (510/580 nm) were obtained throughout the IR injury protocol, and were normalized as described in the methods. **(A):** Signal obtained in the absence of mitoSOX ("no probe" control). **(B):** Signal obtained in the presence of oxidized mitoSOX. 5 min. period of dye loading is indicated by the black bar. **(C):** Signal obtained in the presence of naïve mitoSOX. 5 min. period of dye loading is indicated by the black bar. **(D):** Corrected mitoSOX data, obtained by subtracting the data in panel B from those in panel C, and normalizing the signal for the 5 min. immediately prior to reperfusion, to a value of 1. All traces show means ± SEM, N= 4-6.

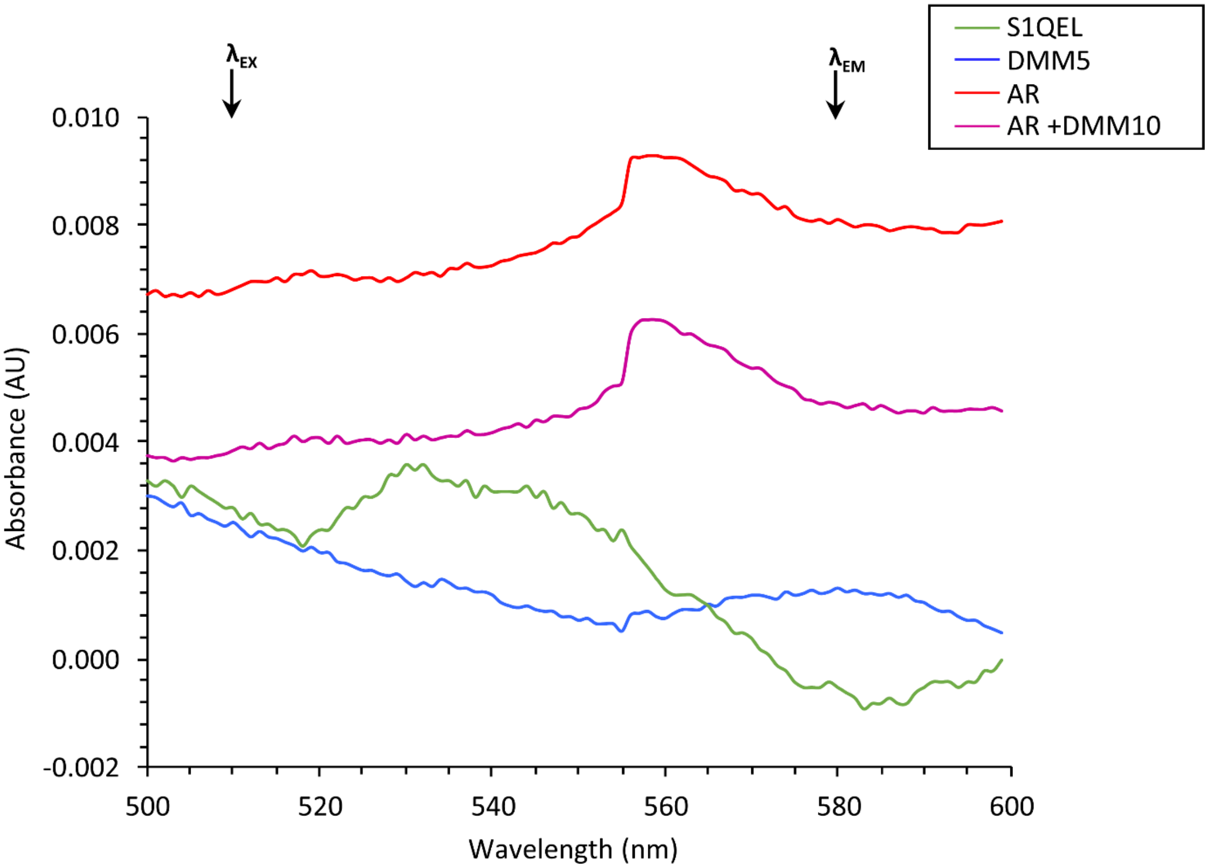


**Supplemental Figure 2. Wavelength Scans for Reagents used in Perfusions.** Absorbance spectra were obtained using a Beckman DU800 UV/Vis spectrophotometer with tungsten and deuterium lamps, quartz cuvets. Graphs show difference spectra (normalized against Krebs-Henseleit buffer) for 1.6 µM S1QEL, 5 mM dimethyl malonate, 10 µM AR-C155858, or 10 µM AR-C155858 plus 10 mM DMM. Colors as per scheme in Figure 1. Excitation and emission wavelengths for mitoSOX are indicated. None of the reagents resulted in a greater than 8 milli OD unit change in absorbance at the required wavelengths.

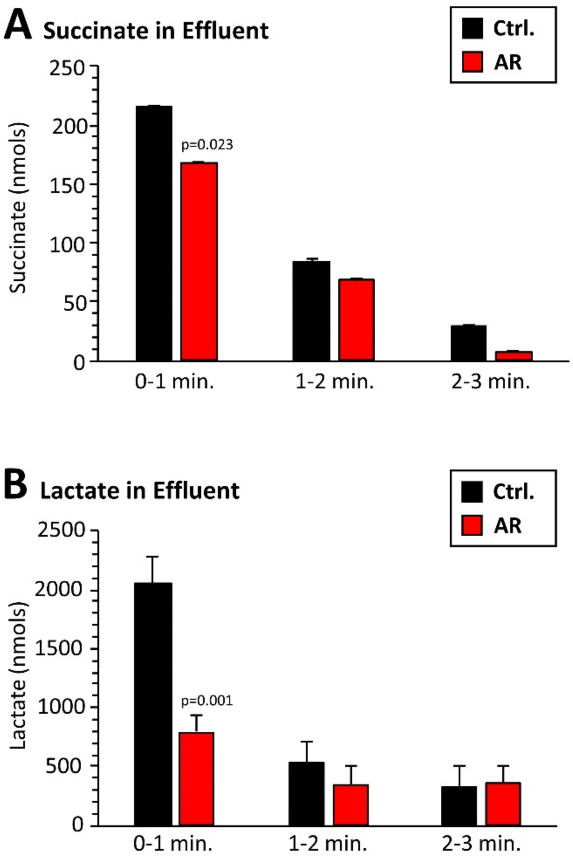


**Supplemental Figure 3. HPLC Quantitation of Metabolites in Cardiac Effluent.** Cardiac perfusates (effluents) were collected in 1 ml aliquots for the first 3 min. of reperfusion, from control hearts or those treated with AR. Perfusion was at a rate of 4 ml/min, so each minute yielded 4 ml of perfusate. Metabolites were analyzed by HPLC with UV/Vis detection as described [1]. Data for **(A):** succinate and **(B):** lactate, are shown as means ± SEM, N=10.

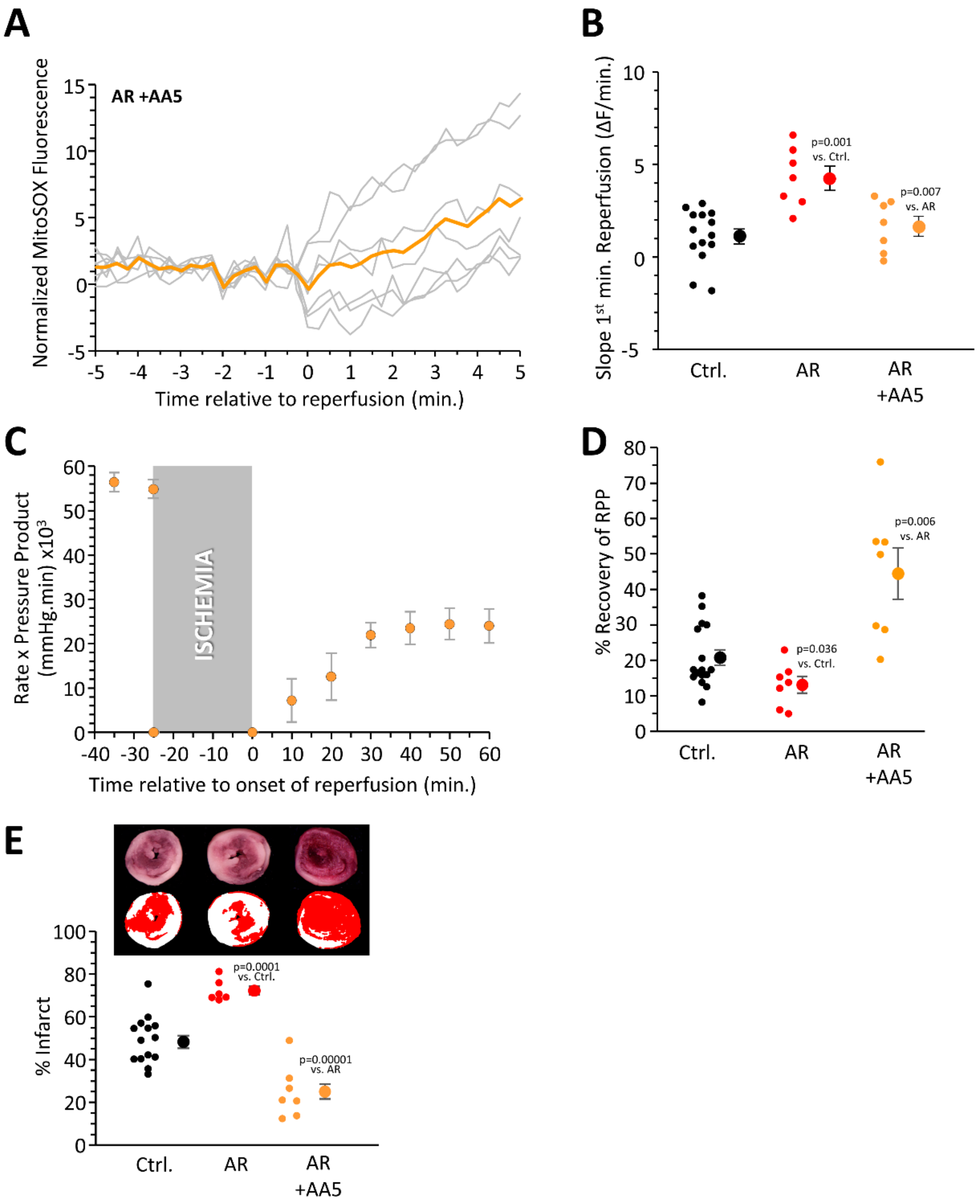


**Supplemental Figure 4. Data for Atpenin A5.** Hearts were treated with AR, or AR plus 100 nM AA5 (see Figure 1 scheme). **(A):** mitoSOX signal upon reperfusion. Gray traces show individual data, with averages shown as bold line. **(B):** Quantitation of mitoSOX slope. Data for control and AR conditions are as per Figure 3F. **(C):** Cardiac function. **(D):** Quantitation of cardiac functional recovery. Data for control and AR conditions are as per Figure 4B. **(E):** Infarct size. Data for control and AR are as per Figure 4C. N for each condition (mitoSOX measurements or cardiac parameters) is indicated by the number of individual data points in panels B and D. Data are means ± SEM. p values (ANOVA followed by unpaired Student's t-test) for differences between groups are denoted.

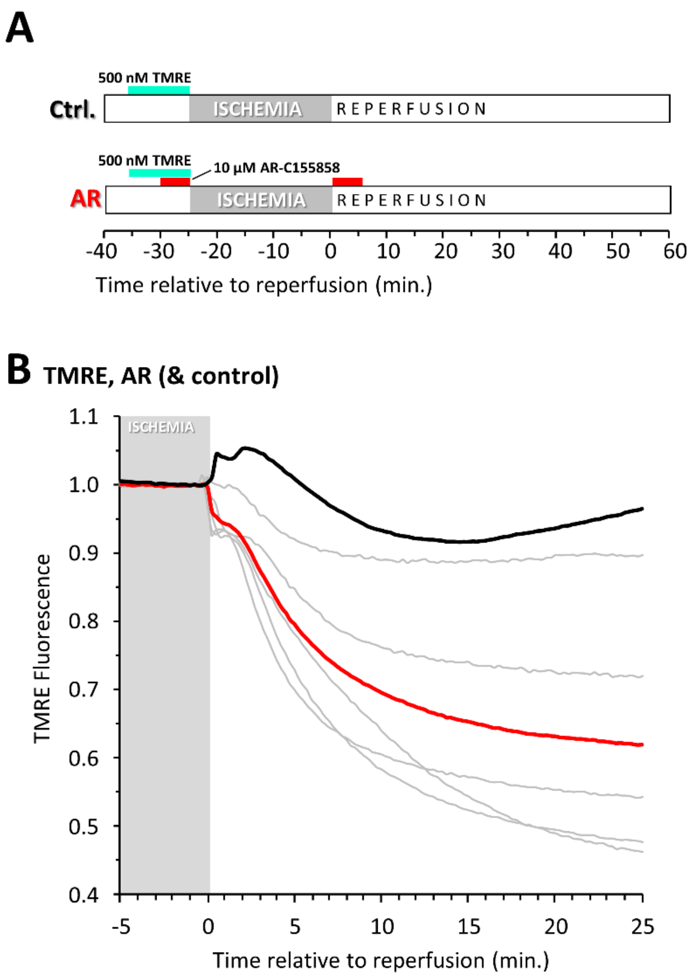


**Supplemental Figure 5. Impact of MCT-1 Inhibition on PT Pore Opening During IR. (A):** Schematic showing perfusion conditions. Where indicated, TMRE ($\Delta\Psi_m$ indicator) and AR-C155858 (MCT-1 inhibitor) were administered at the listed concentrations. **(B):** Normalized TMRE fluorescence during early reperfusion under control or AR condition. The last 5 min. of the ischemic period is indicated. Gray traces show individual data, with the average of the AR condition shown in bold red. Average control data from Figure 7D of the main manuscript are shown in black.
